# Microbial community-scale metabolic modeling predicts personalized short chain fatty acid production profiles in the human gut

**DOI:** 10.1101/2023.02.28.530516

**Authors:** Nick Quinn-Bohmann, Tomasz Wilmanski, Katherine Ramos Sarmiento, Lisa Levy, Johanna W. Lampe, Thomas Gurry, Noa Rappaport, Erin M. Ostrem, Ophelia S. Venturelli, Christian Diener, Sean M. Gibbons

## Abstract

Microbially-derived short chain fatty acids (SCFAs) in the human gut are tightly coupled to host metabolism, immune regulation, and integrity of the intestinal epithelium. However, the production of SCFAs can vary widely between individuals consuming the same diet, with lower levels often associated with disease. A systems-scale mechanistic understanding of this heterogeneity is lacking. We present a microbial community-scale metabolic modeling (MCMM) approach to predict individual-specific SCFA production profiles. We assess the quantitative accuracy of our MCMMs using *in vitro*, *ex vivo*, and *in vivo* data. Next, we show how MCMM SCFA predictions are significantly associated with blood-derived clinical chemistries, including cardiometabolic and immunological health markers, across a large human cohort. Finally, we demonstrate how MCMMs can be leveraged to design personalized dietary, prebiotic, and probiotic interventions that optimize SCFA production in the gut. Our results represent an important advance in engineering gut microbiome functional outputs for precision health and nutrition.

## Introduction

The human gut microbiota serves many functions: maintaining intestinal barrier function, regulating peripheral and systemic inflammation, and breaking down indigestible dietary components and host substrates into a wide range of bioactive compounds ^1,2^. One of the primary mechanisms by which the gut microbiota impacts human health is through the production of small molecules that enter the circulation and are absorbed and transformed by host tissues ^3–5^. Approximately half of the metabolites detected in human blood are significantly associated with cross-sectional variation in gut microbiome composition ^6^.

Short chain fatty acids (SCFAs) are among the most abundant metabolic byproducts produced by the gut microbiota, largely through the fermentation of indigestible dietary fibers and resistant starches, with acetate, propionate and butyrate being the most abundant SCFAs ^7–9^. Deficits in SCFA production, specifically butyrate and propionate, have been repeatedly associated with disease, including inflammatory bowel disease and colorectal cancer ^10–15^. Therefore, SCFA production is a crucial ecosystem service that the gut microbiota provides to its host, with extensive impacts on health ^1,11,16,17^. However, different human gut microbiota provided with identical dietary substrates can show variable SCFA production profiles ^18,19^, and predicting this heterogeneity remains a fundamental challenge to the microbiome field. Measuring SCFA abundances in blood or feces is rarely informative of *in situ* production rates, due to the volatility of SCFAs, cross-feeding among microbes, and the rapid consumption and transformation of these metabolites by the colonic epithelium ^10,20,21^. Furthermore, SCFA production fluxes (i.e., the amount of a metabolite produced over a given period of time) within an individual can vary longitudinally, depending upon dietary inputs and the availability of host substrates ^22^. In order to account for this inter-and intra-individual heterogeneity, we propose the use of microbial community-scale metabolic models (MCMMs), which mechanistically account for metabolic interactions between gut microbes, host substrates, and dietary inputs, to estimate personalized, context-specific SCFA production profiles.

Statistical modeling and machine-learning approaches for predicting metabolic output from the microbiome have shown promising results in recent years. For example, postprandial blood glucose responses can be predicted by machine-learning algorithms trained on large human cohorts ^23,24^. Nevertheless, machine-learning methods are limited by the measurements and interventions represented within the training data ^25^. Mechanistic models like MCMMs, on the other hand, do not rely on training data and can provide causal insights ^21^. Metabolic modeling of individual commensal taxa has been used to predict plasma concentrations of microbially derived metabolites ^26^, but these methods have not been extended to diverse, real-world microbiomes. MCMMs can be constructed using existing knowledge bases, including curated genome-scale metabolic models (GEMs) of individual taxa ^27^. MCMMs are limited by the availability of well-curated GEMs for abundant taxa present within every individual in a population and by information on individual-specific dietary variation. These limitations are further exacerbated in human populations that are generally underrepresented in microbiome research, where our databases are also less representative ^28^. However, as our knowledge bases grow, so too will the power and scope of MCMMs. Overall, MCMMs have the potential to serve as powerful, transparent, knowledge-driven tools for predicting community-specific responses to a wide array of interventions or perturbations.

Here, we demonstrate the utility of MCMMs for the prediction of personalized SCFA production profiles in the context of different dietary, prebiotic, and probiotic inputs. We first validate our modeling platform using diverse synthetic *in vitro* gut microbial communities (N = 1,387) and *ex vivo* stool incubation assays (N = 29). Next, we investigate the relevance of this modeling strategy *in vivo* using data from a 10-week high-fiber dietary intervention cohort (N = 18), where individuals showed a variety of immune responses. We assess the clinical significance of these precision SCFA predictions by looking at associations between predicted SCFA production on an average European diet and a panel of blood-based clinical lab tests in a large human cohort (N = 2,687). Finally, we demonstrate the potential power of MCMMs in designing personalized prebiotic, probiotic, and dietary interventions that optimize predictions for individual-specific butyrate production rates.

## Results

### MCMMs capture SCFA production rates in vitro

Details on the origin and composition of each dataset used in these analyses can be found in the supplement (**Table S1**).

We sought to investigate whether MCMMs can predict production rates of the major SCFAs (i.e., acetate, propionate, and butyrate) under controlled experimental conditions (**Fig. 1**). Growth media, matching the environmental context of each experiment, were constructed and applied as bounds on metabolic import to MCMMs (**Fig. 1A**), which were concurrently constructed by combining manually-curated GEMs from the AGORA database ^29^ using MICOM ^21^, constraining taxon abundances using 16S amplicon or shotgun metagenomic sequencing relative abundance estimates (**Fig. 1B**). Sample-specific metabolic models were then solved using cooperative tradeoff flux balance analysis (ctFBA), a previously-reported two-step quadratic optimization strategy that yields empirically-validated estimates of the steady state growth rates and metabolic uptake and secretion fluxes for each taxon in the model ^21^ (**Fig. 1C**, see Materials and Methods). Models constructed from 16S amplicon sequencing data were summarized at the genus level, which was the finest level of phylogenetic resolution that the data allowed for. When shotgun metagenomic sequencing data were available, models were constructed at the species level. Models constructed from both 16S and shotgun metagenomic data at the species and genus levels showed highly consistent results (**Fig. S1**). Measured SCFA production profiles from synthetic *in vitro* community and stool *ex vivo* experiments (**Fig. 1D**) were compared to paired SCFA flux predictions from MCMMs to validate the accuracy of the models.

**Figure 1.**
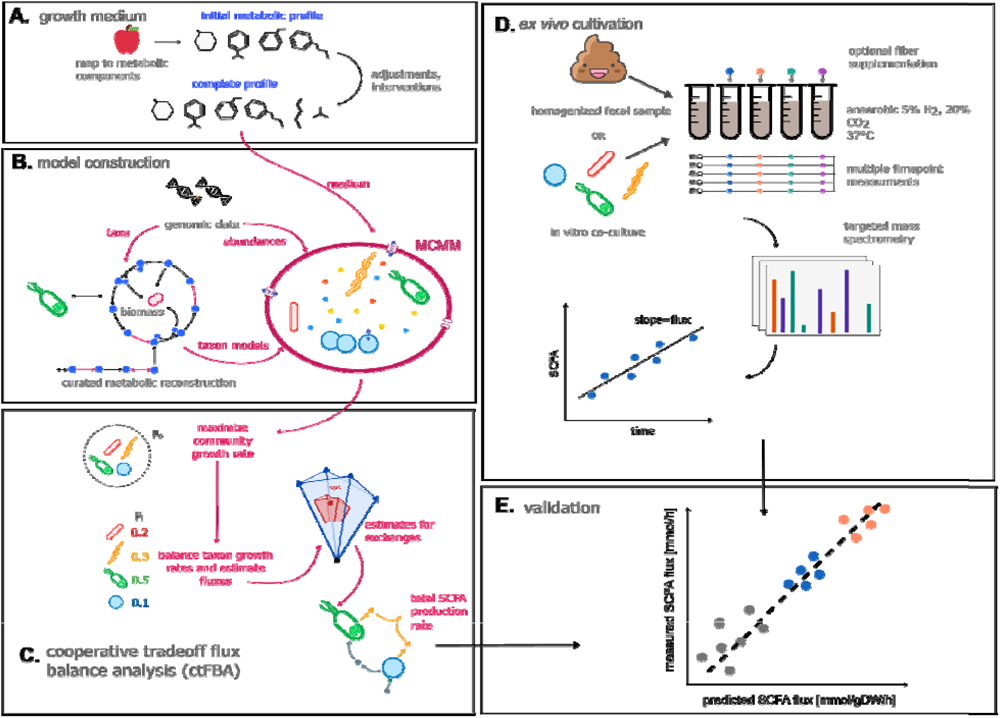
Microbial community-scale metabolic models (MCMMs) predict personalized SCFA production profiles. Schematic of our workflow for validating MCMM-based personalized predictions for SCFA production. **(A)** Prior to modeling, an *in silico* medium is constructed, containing a matched diet mapped to its constituent metabolic components. The medium is depleted in compounds absorbed by the host in the small intestine and augmented with other host-supplied compounds, in addition to adding a minimal set of metabolites required for growth. **(B)** MCMMs are constructed, combining abundance and taxonomic data with pre-curated GEMs into a community model. **(C)** Growth in the MCMM is simulated through cooperative tradeoff flux balance analysis (ctFBA), yielding predicted growth rates and SCFA production fluxes. **(D)** To validate predicted levels of SCFA production fluxes, measured production fluxes are collected from *in vitro* communities of human gut commensals and fecal samples cultured anaerobically *ex vivo* at 37°C over time. **(E)** Predicted and measured SCFA production fluxes are compared to assess the accuracy of the model.

First, we looked at published data from synthetically constructed communities of bacterial commensals isolated from the human gut ^30^. This data set included endpoint measurements of relative microbial abundances, derived from 16S amplicon sequencing, measured endpoint butyrate concentrations, and the overall optical density for each of 1,387 independent co-cultures (**Fig. 2A**). Cultures varied in richness from 1-25 strains. MCMMs were constructed for each co-culture as described above, simulating growth of each of the models using a defined medium mapped to a database of metabolic constituents, matching the composition of the medium used in the *in vitro* experiments (see Materials and Methods). Model-predicted butyrate fluxes were compared with calculated butyrate production rates (endpoint butyrate divided by culturing time, assuming no butyrate at the start of growth, normalized to total biomass using OD600), stratifying results into low richness (1-5 genera) and high richness (10-25 genera) communities. Model predictions for butyrate production fluxes were significantly correlated with measured butyrate production fluxes across all communities (Pearson’s correlation; Low Richness: r = 0.17, p < 0.001; High Richness: r = 0.53, p < 0.001), but the predictions were more accurate in the higher richness communities (**Fig. 2B-C**).

**Figure 2.**
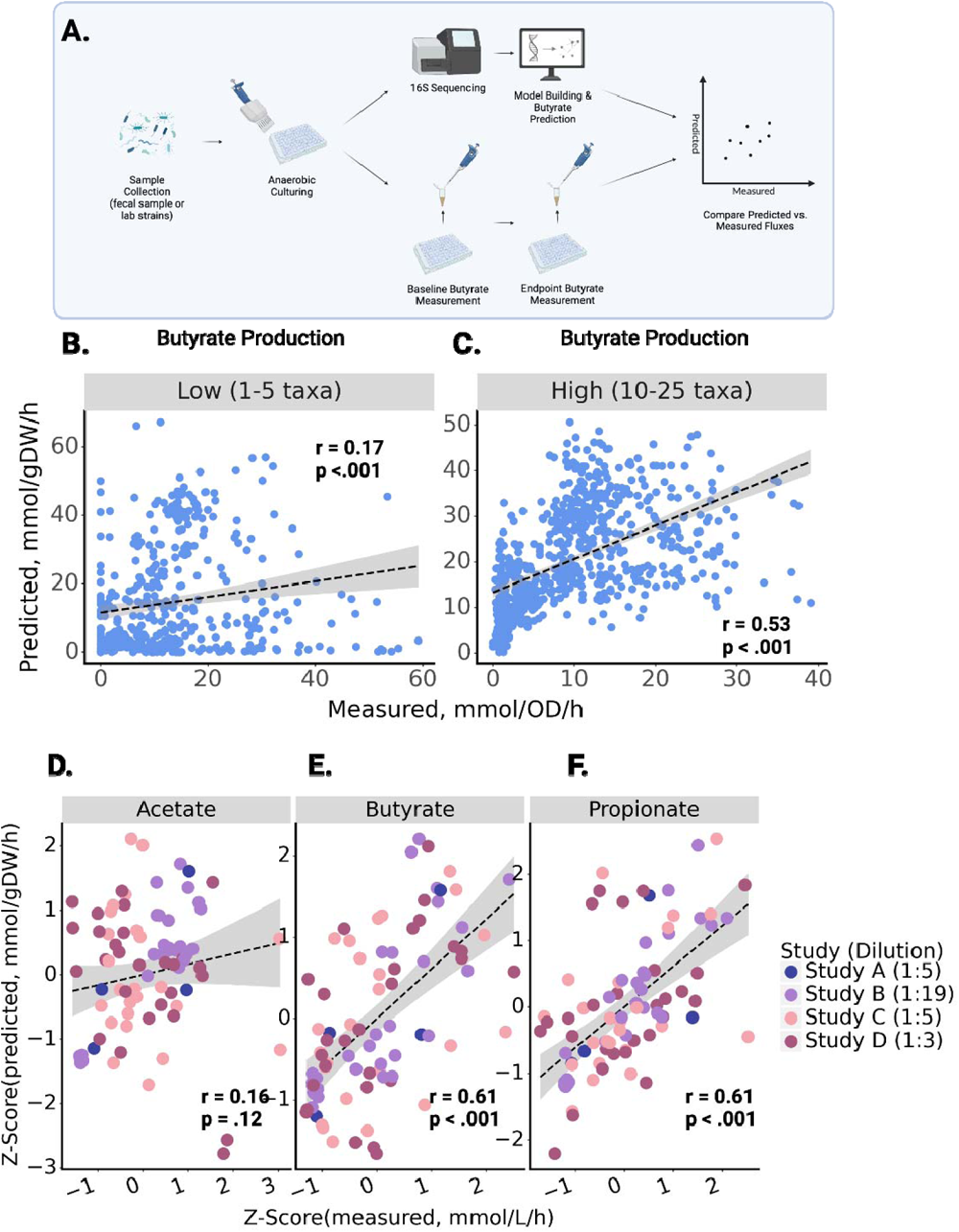
Relationship between predicted and measured butyrate production rates in *in vitro* and *ex vivo* co-cultures. Butyrate production flux predictions from MCMMs are shown on the y-axes and measured values are shown on the x-axes, along with R^2^ and p-values from a Pearson’s correlation **(A)** *In vitro* or *ex vivo* communities were cultured anaerobically. Endpoint butyrate concentration was used to calculate production flux and compared with MCMM-predicted flux. **(B)** Predicted and measured butyrate fluxes in models of low richness synthetic communities (1-5 genera per model, N = 882). **(C)** Predicted and measured butyrate fluxes in models of high richness synthetic communities (10-25 genera, N = 697). **(D-F)** Z-scored predicted and measured fluxes for acetate, butyrate and propionate, across four independent *ex vivo* studies. The label in the figure legend indicates the final dilution level of cultures in each study (dilution = 1:x). In (B-F) the dashed line denotes a linear model fit to the data, with the surrounding shaded region indicating the 95% confidence interval.

Next, we compared MCMM predictions to anaerobic *ex vivo* incubations of human stool samples from a small number of individuals (N = 29), cultured after supplementation with sterile PBS buffer or with different dietary fibers across four independent studies. Study A contained samples from two donors cultured for 7 hours with a final dilution of 1:5, Study B ^18^ contained samples from 10 donors cultured for 24 hours diluted 1:19, Study C contained samples from 8 donors cultured for 4 hours diluted 1:5, and Study D contained samples from 9 donors cultured for 6 hours diluted 1:3. Fecal *ex vivo* assays allow for the direct measurement of bacterial SCFA production fluxes without interference from the host. For all three studies, *ex vivo* incubations were performed by homogenizing fecal material in sterile buffer under anaerobic conditions, adding control or fiber interventions to replicate fecal slurries, and measuring the resulting SCFA production rates *in vitro* at 37°C (see Materials and Methods). Metagenomic (Studies A, C and D) or 16S amplicon (Study B) sequencing data from these *ex vivo* cultures were used to construct MCMMs, using relative abundances as a proxy for relative biomass for each bacterial taxon (see Materials and Methods). MCMMs were simulated using a diluted standardized European diet (i.e., to approximate residual dietary substrates still present in the stool slurry), with or without specific fiber amendments, matching the experimental treatments (see Material and Methods). Within studies, the divergence in measured SCFA production between control samples and fiber-treated samples seemed to be highly dependent upon the final dilution of the *ex vivo* cultures (**Fig. S2**). This was accounted for by matching the dilution of residual fiber (starch, cellulose and dextrin) in the medium used for growth simulation to the corresponding study. For instance, Study A was diluted 1:5, so the residual fiber in the medium used to simulate growth for these samples was diluted by a factor of 5. The resulting SCFA flux predictions were then compared to the measured fluxes. MCMM fluxes are given in units of mmol/gDW/h, while measured production fluxes are given in mmol/L/h. Without knowledge of the live-cell biomass within the fecal homogenates, it was not possible to normalize the units across the two axes, but the predicted and measured values were expected to be proportional. To overcome study-specific differences in protocols and allow for comparison of results across studies, we Z-scored both measured and predicted SCFA production fluxes within each data set (**Fig. 2D-F**). We observed a similar degree of agreement between MCMM-predicted and measured production fluxes for butyrate and propionate across all four *ex vivo* data sets (**Fig. 2E-F**). The model was notably less capable of accurately predicting differences in acetate production between individuals, with no significant association seen (**Fig. 2-3**). Significant agreement was observed between measured and predicted production fluxes of butyrate and propionate within each individual data set (r =0.41-0.97, Pearson test, p<0.05) with the exception of propionate in Study A, which had a very limited sample size (N = 2) (**Fig. 3E-L**). Notably, the correlation coefficient (Pearson r) for these associations was similar to that seen in the high-richness *in vitro* cultures (**Fig. 2C**). As previously seen, the prediction of acetate was worse, most notably in studies C and D, where no significant prediction was observed. In studies A and B, acetate production was more readily predicted, likely due to a strong treatment-effect (**Fig. 3A-D**). Within treatment groups, similar correlations were observed, though statistical power was severely limited by the smaller sample sizes (**Table S2**). Predictions from models built with shotgun metagenomic sequencing data showed slightly better results when constructed at the species level, as compared to building at the genus level (**Fig. S3**). To test whether SCFA production was impacted by sample diversity, we compared measured butyrate and propionate against Shannon index for each sample in each study (**Fig. S4**). A weak significant signal was seen in only one of the four studies (Study D). In summary, we observed agreement between MCMM predicted and measured *in vitro* production rates of butyrate and propionate in the presence or absence of fiber supplementation, with better agreement in more diverse communities and over longer experimental incubation times (**Fig. 2**-**3**). As acetate was not well predicted by the MCMMs (i.e., acetate was not strongly coupled to biomass production, and predictions could vary widely for the same biomass optimum), we focused our downstream predictions and analyses on the SCFAs butyrate and propionate.

**Figure 3.**
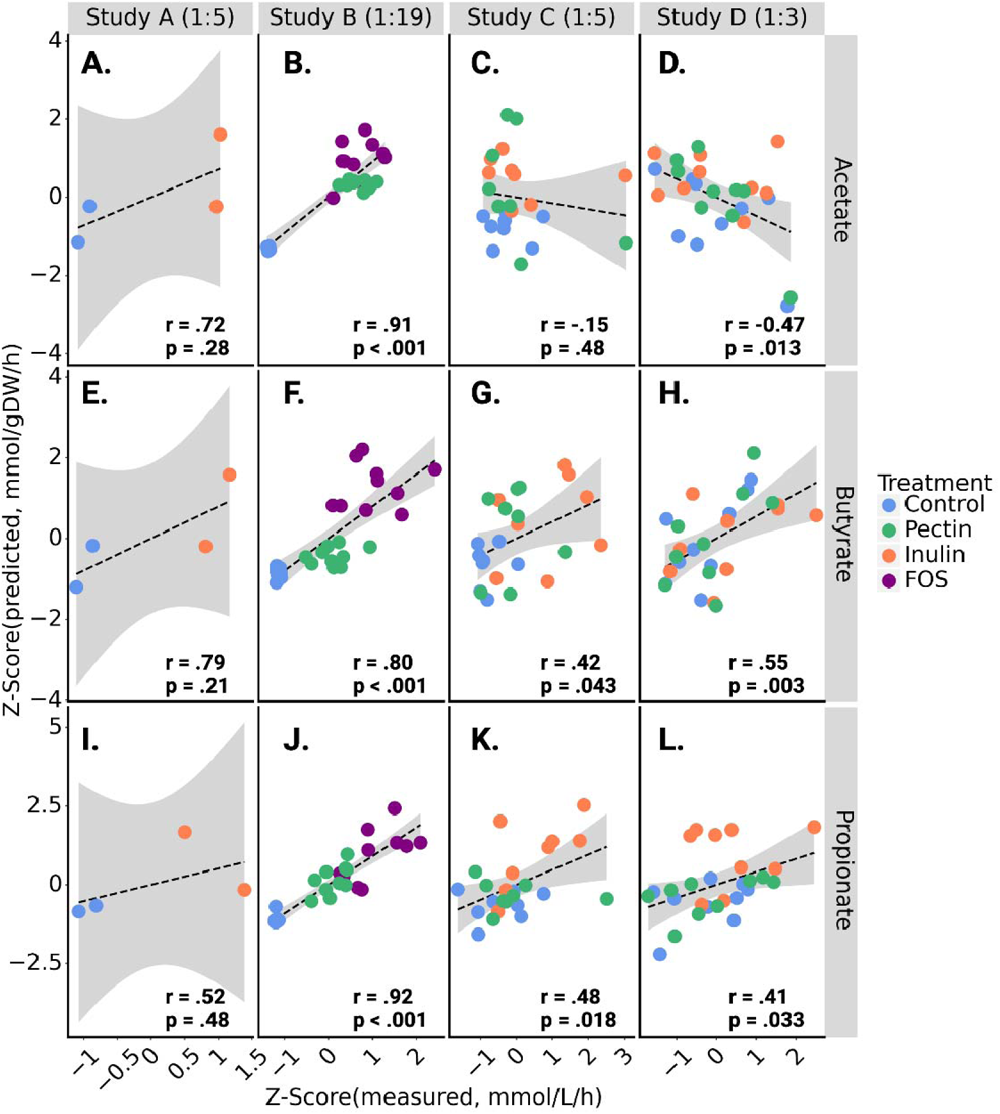
Human stool *ex vivo* assays show quantitative agreement between measured and predicted SCFA production fluxes within and across fiber treatment groups. Z-scored SCFA production flux predictions from MCMMs are shown on the y-axes and Z-scored measured values are shown on the x-axes. Pearson’s r and associated p-value are calculated for all points in a given plot. Color encoding indicates the specific fiber treatment given to each sample. The dashed line denotes a linear regression line and the gray area denotes the 95% confidence interval of the regression. Residual fiber in the media used to simulate growth of each study was scaled according to the dilution factor, shown next to the study name in each column **(A-D)** Z-scored predictions compared with z-scored measurements of acetate production across all four studies. **(E-H)** Z-scored predictions compared with z-scored measurements of butyrate production across all four studies. **(I-L)** Z-scored predictions compared with z-scored measurements of propionate production across all four studies.

### MCMM predictions correspond with variable immunological responses to a 10-week high-fiber dietary intervention

We next investigated whether MCMM-predicted SCFA production rates could be leveraged to help explain inter-individual differences in phenotypic response following a dietary intervention. Specifically, we looked at data from 18 individuals who were placed on a high-fiber diet for ten weeks ^31^. These individuals fell into three distinct immunological response groups: one in which high inflammation was observed over the course of the intervention (high-inflammation group), and two other distinct response groups that both exhibited lower levels of inflammation (low-inflammation groups I and II; **Fig. 4A**). We hypothesized that these immune response groups could be explained, in part, by differences in MCMM-predicted production rates of anti-inflammatory SCFAs. Using 16S amplicon sequencing data from seven time points collected from each of these 18 individuals over the 10-week intervention, we built MCMMs for each study participant at each time point. Growth was then simulated for each model using a standardized high-fiber diet, rich in resistant starch (see Material and Methods). Throughout the study, a trend of decreasing propionate production was observed in high-inflammation individuals (r = 0.39, Pearson test, p = 0.019), showing less production as the intervention went on, despite the high fiber content of the diets consumed by participants (**Fig. 4B**). Individuals in the high-inflammation group showed significantly lower predicted propionate production, on average, compared to the individuals in each of the low-inflammation groups (High vs. Low I: 131.9 ± 5.8 vs 158.1 ± 5.7 mmol/(gDW h), Mann-Whitney p = 0.0053; High vs. Low II: 131.9 ± 5.8 vs 163.08.3 ± 6.5 mmol/(gDW h), Mann-Whitney p = 0.0017; **Fig. 4C**). Butyrate showed no such significant effects across immune response groups (**Fig. 4D, 4E**). To investigate whether sample alpha-diversity was sufficient to explain the differences between the immune response groups, we calculated the alpha diversity for each sample at each timepoint during the study. Across all seven time points tested, only one significant difference in alpha diversity was seen, between the two low inflammation groups at time point 2 (Mann-Whitney U-test, p < 0.05), leading us to determine that differences in SCFA production throughout the intervention were not the result of differences in diversity. (**Fig. S4**).

**Figure 4.**
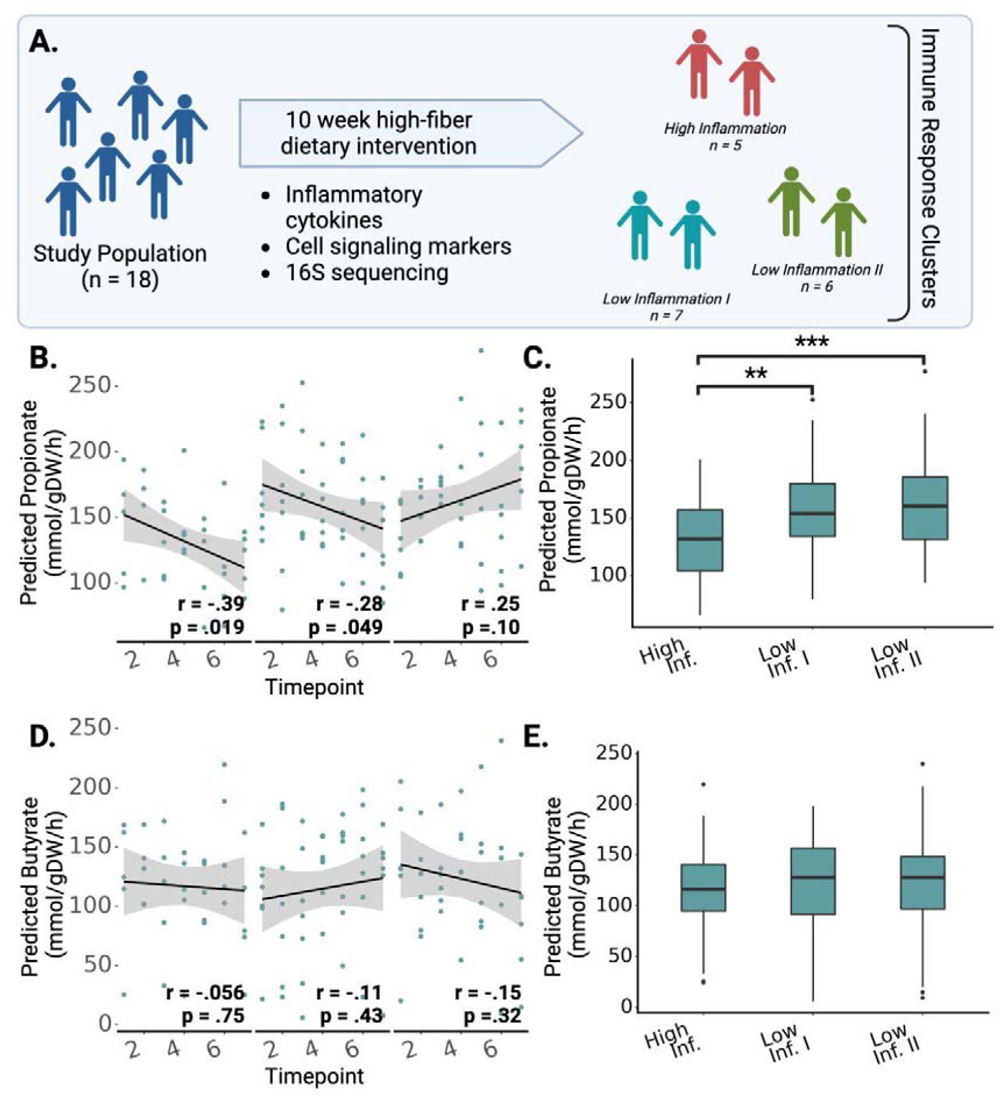
Predicted SCFA production profiles were associated with variable immune response groups following a high-fiber dietary intervention. **(A)** Summary of the study from Wastyk et al.^31^, where a cohort of 18 individuals participated in a 10-week high-fiber dietary intervention. Immune profiling based on circulating inflammatory cytokines and immune cells clustered individuals into three groups: two low-inflammation groups and one high-inflammation group. **(B)** Total predicted propionate production at each timepoint across the three immune-response groups identified in the original study. **(C)** Average predicted propionate production rates, stratified by immune response group **(D)** Total predicted butyrate production at each timepoint across the three immune-response groups identified in the original study. **(E)** Average predicted butyrate production rates, stratified by immune response group. In (B-E) stars denote significance under a Mann-Whitney U-test, * = p<0.05, ** = p< 0.01, *** = p<0.001.

### MCMM-predicted SCFA profiles are associated with a wide range of blood-based clinical markers

To further evaluate the clinical relevance of personalized MCMMs, we generated SCFA production rate predictions from stool 16S amplicon sequencing data for 2,687 individuals in a deeply phenotyped, generally-healthy cohort from the West Coast of the United States (i.e., the Arivale cohort) ^32^. Baseline MCMMs were built for each individual assuming the same dietary input (i.e., an average European diet) in order to compare SCFA production rate differences, independent of background dietary variation. MCMM-predicted SCFA fluxes were then regressed against a panel of 128 clinical chemistries and health metrics collected from each individual, adjusting for a standard set of common covariates (i.e., age, sex, and microbiome sequencing vendor; **Fig. 5A**). After FDR correction, 20 markers were significantly associated with the predicted production rate of butyrate (**Fig. 5B**). Predicted butyrate production showed significant positive associations with only 3 markers, including the health-associated hormone adiponectin, and significant negative associations with 17 markers linked to disease, including C-reactive protein (CRP), low-density lipoprotein (LDL), and blood pressure (mean arterial pressure; P < 0.05, FDR-corrected t-test). Propionate showed no significant associations after covariate adjustment and multiple comparison correction (**Fig. 5B**). Total combined propionate and butyrate production was significantly associated with 16 clinical markers, all overlapping with those associated with butyrate. Predicted butyrate production was significantly negatively associated with BMI (β = -0.10, t-test, p < 0.001), while propionate was not (**Fig. 5** **C-D**). Covariate-adjusted p-values and beta coefficients for all clinical markers included in the analysis can be found in the supplementary material (**Table S3**).

**Figure 5.**
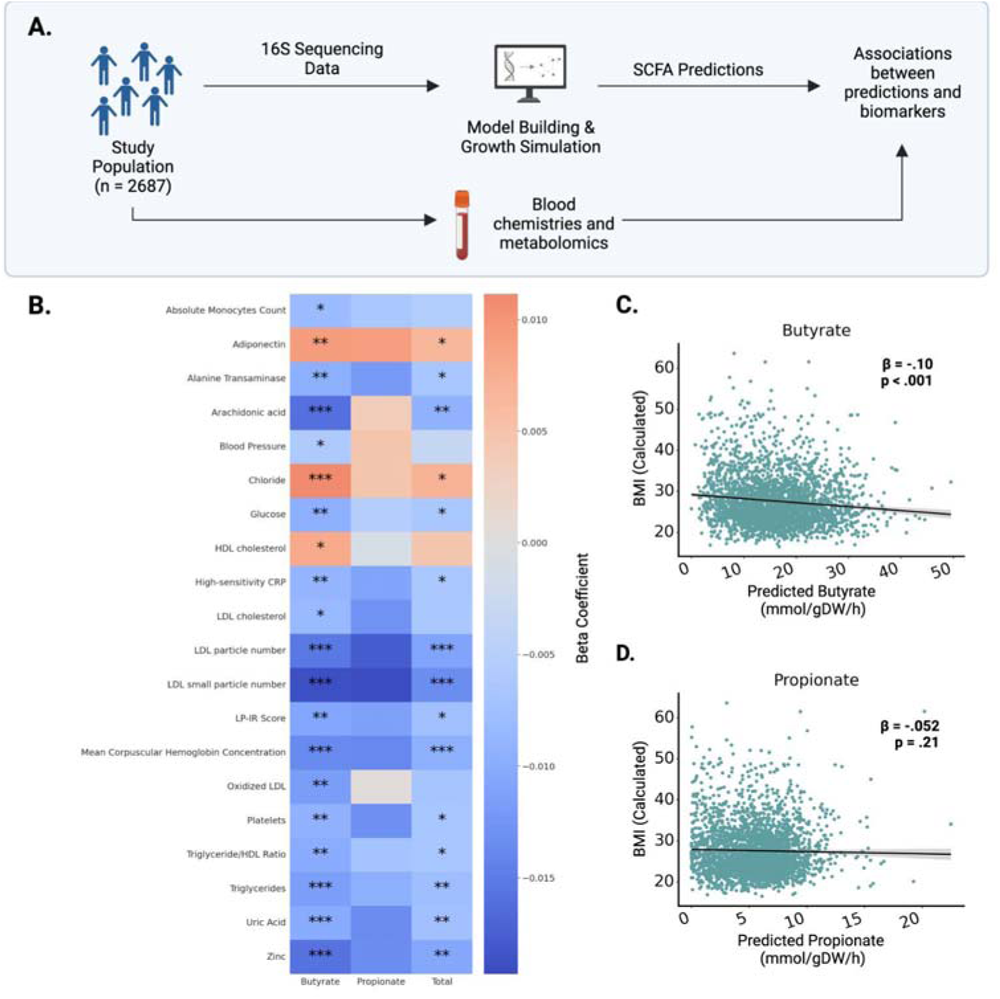
SCFA flux predictions are significantly associated with blood-derived clinical markers. **(A)** MCMMs were constructed for 2,687 Arivale participants, assuming an average European diet, to predict SCFA production profiles. SCFA predictions were regressed against a set of 128 blood-based clinical labs and health markers, with sex, age, and sequencing vendor as covariates in the regressions. **(B)** Heatmap showing the 20 significant associations (FDR-corrected t-test p<0.05) between measured blood markers and predicted SCFA production rates. **(C-D)** Predictions for butyrate were significantly correlated with reported BMI measurements for respective participants, but not for propionate. Each dot denotes an individual model reconstructed for a single sample in the Arivale study (N = 2,687). The black line denotes a linear regression line and the gray area denotes the 95% confidence interval of the regression. β-coefficients were calculated from multiple regression accounting for age, sex and microbiome sequencing vendor.

### Leveraging MCMMs to design precision dietary, prebiotic, and probiotic interventions

As a proof-of-concept for *in silico* engineering of the metabolic outputs of the human gut microbiome, we screened a set of potential interventions designed to increase SCFA production for individuals from the Arivale cohort (**Fig. 6A**). MCMMs were built using two different dietary contexts: an average European diet, and a vegan, high-fiber diet rich in resistant starch (see Material and Methods). As expected, models grown on a high-fiber diet showed higher average predicted butyrate production: 87.78 ± 0.67 mmol/(gDW h) vs 16.29 ± 0.13 mmol/(gDW h), t-test, t = 104.3, p < 0.001 (**Fig. 6B**). However, this increase in butyrate production between the European and high-fiber diets was not uniform across individuals. On the high-fiber diet, some individual gut microbiota compositions showed very large increases in butyrate production, some showed little-to-no change, and a small subset of samples actually showed a decrease in butyrate production, relative to the European diet. We identified a set of ‘non-responders’ (N = 9) who produced less than 15 of butyrate on the European diet and showed an increase in butyrate production of less than 20% on the high-fiber diet (**Fig. 6C**). We also identified a set of ‘regressors’ (N = 7) who showed decreased butyrate production on the high-fiber diet when compared to the European diet (**Fig. 6D**). We then simulated a handful of simple prebiotic and probiotic interventions across these individuals, to identify optimal combinatorial interventions for each individual (**Fig. 6C-E**). MCMMs for each subset of individuals were simulated with prebiotic and probiotic interventions in the context of either the European or the high-fiber diet. Specifically, diets were supplemented with the dietary fiber inulin, with the dietary fiber pectin, or with a simulated probiotic intervention that consisted of introducing 10% relative abundance of the butyrate-producing genus *Faecalibacterium* to the MCMM. In general, optimal combinatorial interventions significantly increased the population-level butyrate production well above either dietary intervention alone (**Fig. 6C-D**).

**Figure 6.**
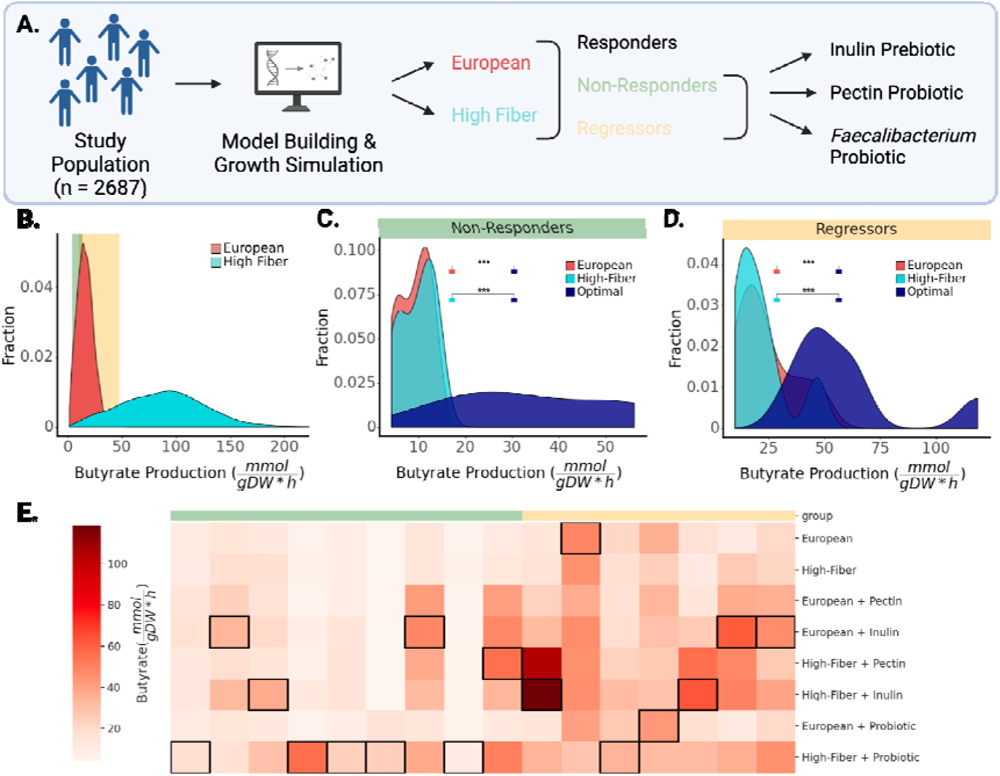
Microbial MCMMs can be used to design and select personalized prebiotic, probiotic, and dietary interventions aimed at optimizing SCFA production profiles. **(A)** MCMMs built from the Arivale cohort (N = 2,687) were used to test personalized responses to dietary interventions. Personalized models were simulated on an average European diet, as well as on a high-fiber diet, and divided into responders, non-responders, and regressors, based on the changes in predicted butyrate production in response to increasing dietary fiber. Non-responders were defined as individuals who produced less than 15 of butyrate on the European diet and showed an increase of less than 20% in butyrate production on the high-fiber diet. Regressors were defined as individuals who showed a decline in butyrate production on the high-fiber diet when compared to the European diet. Single-fiber and probiotic interventions were applied to non-responders and regressors. **(B)** Distribution of butyrate production rates on two different diets simulated for all participants in the study. Butyrate production ranges that contain non-responders (N = 9) and regressors (N = 7) are highlighted in green and yellow shaded areas, respectively. **(C)** Distributions of butyrate production rates for the non-responder group (N = 9). The optimal intervention resulting in the highest butyrate production is shown in blue. **(D)** Butyrate production rates for the regressor group (N = 7). The optimal intervention that resulted in the highest butyrate production is shown in blue. **(E)** Heatmap of butyrate production rates across simulated interventions for the individuals in the non-responder and regressor groups. Rows denotes specific interventions, columns denote individuals in the response groups (N = 16). Cell shading (white-to-red) denotes butyrate production rate. Added interventions tested on both non-responders and regressors included probiotic supplementation (inulin or pectin) as well as prebiotic supplementation (5% relative abundance *Faecalibacterium*). The most successful intervention for each individual is denoted by a black border around that cell in the corresponding column.

For 15/16 individuals in the regressors or non-responders groups, supplementation of the background diet with a specific prebiotic or probiotic increased the butyrate production rate (**Fig. 6C-E**). For both regressors and non-responders, the optimal intervention showed substantial increases over the standard European diet (+290±80% for non-responders; +239±102% for regressors). The exact intervention that yielded the highest butyrate production for any given individual across both populations varied widely (**Fig. 6E**). For example, the probiotic intervention was more successful in raising predictions for butyrate production in non-responders than it was in regressors (**Fig. 6E**). Overall, no single combinatorial intervention was optimal for every individual in the population.

## Discussion

The objective of this study was to experimentally validate personalized MCMM SCFA predictions. Predictions of butyrate production in synthetically constructed *in vitro* co-cultures showed significant agreement between measured and predicted butyrate fluxes (**Fig. 2**), a first step toward validation. Interestingly, better agreement was observed in richer communities, indicating increased model complexity benefitted predictions. Decreasing accuracy of butyrate predictions as community richness declined may reflect a limitation of building models at the genus-level, as reconstructions contain a summarized aggregation of the metabolic capability of the genus as a whole, without species-or strain-level resolution. Furthermore, we are leveraging database models, which do not reflect the exact strains present in a given sample. Consequently, pathways included in the metabolic models are not a perfect match to the reality of what is present in a sample. In high richness models, predictions of SCFAs became more accurate, suggesting this mismatch gets averaged out as species richness increases, likely due to functional redundancies across organisms that can mask the inaccuracies of any single taxon model. Alternatively, there could be some unknown biological reason for why SCFA production is less variable in higher richness communities, which would affect our ability to make accurate MCMM predictions. Overall, the observed increase in accuracy with community diversity benefits modeling of real-world microbiomes, which are often more species-rich than synthetic *in vitro* communities ^33,34^. As our model databases grow to better-reflect the metabolic diversity of real-world ecosystems, we expect MCMMs to become more and more accurate, independent of community diversity.

Further validation of MCMM predictions was observed from *ex vivo* anaerobic fecal incubations. We saw good agreement between SCFA flux predictions and measurements, especially for butyrate and propionate, across four independent studies (**Fig. 3**). Acetate is known to act as an overflow metabolite ^35,36^, with a wide range of possible fluxes for a given biomass optimum, so it is perhaps not surprising that the predictions for this metabolite tended to be less accurate across studies and within treatment groups. Butyrate and propionate, however, showed a narrower range of possible fluxes for a given biomass optimum, suggesting that the production of these molecules is more strongly coupled to biomass production. The dilution level of the *ex vivo* stool incubations had a sizable effect on the results, where the *in vitro* prebiotic treatment effect was dampened in less dilute fecal homogenates, likely due to the presence of residual dietary fibers in stool. The more accurate predictions of acetate production in the more dilute fecal homogenates is likely due to the fact that total SCFA production was more strongly coupled to *in vitro* prebiotic treatment in these samples. Accounting for this dilution factor in the construction of the *in silico* media improves predictions and returns more accurate results for butyrate and propionate production.

We were interested in seeing how 16S-and metagenomic-based models compared at a similar taxonomic level, and how genus and species level predictions compared, in order to assess how applicable our modeling strategy could be to different data types. Using paired 16S and shotgun metagenomic sequencing data from Study C, we saw strong agreement between models constructed at the genus level for both 16S and metagenomic data (**Fig. S1**). Furthermore, we saw robust agreement between predictions at the genus and species levels across metagenomic data sets (**Fig. S5**). Interestingly, predictions from Studies A, C and D showed marginally better agreement with measured values when constructed at the species level vs. the genus level, indicating that higher specificity in model construction is desirable when possible (**Fig. S5**). Across the *in vitro* and *ex vivo* studies, our results strongly support the use of MCMMs for predicting personalized butyrate and propionate production rates in response to prebiotic, probiotic, and dietary interventions.

*In vivo* validation via direct measurement of SCFA production is not easily accomplished, due to the rapid consumption of these metabolites by the colonic epithelium and noisy measurements in either stool or serum ^37^ ^38^. However, higher SCFA production rates are known to influence the phenotype of the host in a number of ways, including a reduction in systemic inflammation and improvements in cardiometabolic health ^17,22,39,40^. Wastyk et al. found that among 18 individuals given a 10-week high fiber dietary intervention, one third showed an increase in inflammation over the course of the intervention and two thirds showed a decline in systemic markers of inflammation ^31^. In the original paper, there was no clear mechanism for explaining these variable immune response groups ^31^. We found that propionate production, a strong inhibitor of inflammation through activation of FFA2 and FFA3^41,42^, was predicted to be significantly lower in individuals who showed the high inflammation response (**Fig. 4B-C**) ^31^. While it is impossible to say whether or not our propionate flux predictions are underlying these dietary response phenotypes, the observed immune response groups and propionate production fluxes could not be explained by differences in alpha-diversity between groups (**Fig. S4**). We also had access to blood-based clinical labs and microbiome data for a cohort of 2,687 Americans. We built MCMMs for this cohort, assuming a standard European diet, and predicted butyrate and propionate production. We found that butyrate was negatively associated with systemic inflammation, LDL cholesterol, and insulin resistance, blood pressure, and BMI (**Fig. 5**). These results are consistent with what is known about how butyrate is protective against inflammation, cardiovascular disease, obesity, and metabolic syndrome ^17,22,39,40,43^ (**Fig. 5B**), and they provide us with further confidence in the predictive power of our MCMM approach. Dietary interventions have long been known to elicit variable responses, but a mechanistic framework for predicting this microbiome-mediated heterogeneity has not been available until now.

Given this set of promising associations between SCFA predictions and host phenotypic variation, we sought to demonstrate the potential for leveraging MCMMs for designing precision prebiotic, probiotic, and dietary interventions. Using the Arivale cohort, we identified two classes of individuals that responded differently to an *in silico* high-fiber dietary intervention: non-responders and regressors (**Fig. 6**). We designed combinatorial interventions that added either a prebiotic or a probiotic to the background diets, to see if we could rescue these non-responder and regressor phenotypes. We found significant heterogeneity in which combinatorial intervention was optimal across individuals from each of these response groups (**Fig. 6E**). Given that the non-responders had low baseline levels of butyrate production to begin with and did not respond to a high-fiber diet, this result underscores the importance of personalized predictions for those who tend not to respond well to population-scale interventions. These results also suggest that butyrate production in some individuals is limited by composition of the microbiota, indicating that probiotic interventions would be necessary to induce meaningful increases in production.

This study had several limitations that should be considered. First, we were limited by the availability of high-quality fluxomic data sets for model validation. For example, we had limited sample sizes in the *ex vivo* fecal studies presented above, due to the cost and difficulty of generating these kinds of data for larger cohorts. Additionally, the human cohort data presented here only provided indirect support for our MCMM predictions (**Figs. 4-5**). Second, predictions are dependent on the availability of GEMs. Obtaining large numbers of GEMs that faithfully recapitulate the full metabolic capacities of each organism in a sample is a challenging task. We used the publicly available AGORA model database ^29^. While AGORA models have gone through some degree of manual curation, many of these models are not fully validated and have been shown to include infeasible and missing reactions ^44^. Nevertheless, these GEMs appear to work well in the context of butyrate and propionate flux predictions. SCFA production pathways are fairly phylogenetically conserved and adjacent to central metabolism, so we might expect these reactions to be robust to strain-or species-level variation and variation in model quality. However, predictions for metabolites that are peripheral to central metabolism will likely be much noisier in the absence of well-curated models that closely match the organisms within a given sample. Third, model building is dependent on accurate taxonomic assignment of sequencing reads. For 16S amplicon sequencing, reads can only be confidently assigned at the genus level, limiting the specificity of a model to the genera present in the original samples. However, as model databases grow and shotgun metagenomic sequencing becomes more common, we anticipate this limitation will be resolved. Finally, the lack of individual-specific dietary constraints limits the accuracy of our predictions. For *ex vivo* fecal fermentations, as well as *in vivo* analysis, participant dietary information was not available, and so a standard European diet was used across all models. Detailed knowledge of dietary intake should increase the accuracy of MCMM predictions. Despite these limitations, MCMMs were able to explain 25-35% of the variance in butyrate and propionate production across individuals, and we expect that advances in model curation, pathway annotation, and personalized dietary constraints will only improve upon the accuracy of this approach over time.

## Conclusion

Here we present an approach for the rational prediction of personalized SCFA production rates from the human gut microbiome, validated using *in vitro, ex vivo* and *in vivo* experimental data. Additional analysis demonstrated a clear relationship between SCFA predictions and physiological responses in the host, including lower inflammation and improved cardiometabolic health. SCFA predictions were also significantly associated with variable immune responses to a high fiber dietary intervention. Finally, we showed how MCMMs could be used to rapidly design and test combinatorial prebiotic, probiotic and dietary interventions *in silico* for a large human population. Personalized prediction of SCFA production profiles from human gut MCMMs represents an important technological step forward in leveraging computational systems biology for precision nutrition. Mechanistic modeling allowed us to translate the ecological composition of the gut microbiome into concrete, individual-specific metabolic outputs, in response to particular interventions ^45,46^. MCMMs are transparent models that do not require training data, with clear causal and mechanistic explanations behind each prediction. The clinical relevance of these predictions is evident, due to the widespread physiological effects of SCFAs on the human body ^47,48^. A rational framework for engineering the production or consumption rates of these metabolites has broad potential applications in precision nutrition and personalized healthcare.

## Materials and Methods

### In vitro culturing

Culturing of the synthetically assembled gut microbial communities is described in Clark et al., 2021 ^30^. Culturing of *ex vivo* samples in Study A was done using the methodology described below. Culturing of *ex vivo* samples in Study B is described in Cantu-Jungles et al., 2021^18^. Culturing of *ex vivo* samples in Study C was conducted by co-author Dr. Thomas Gurry, using the methodology described below.

### In vitro culturing of fecal-derived microbial communities (Study A)

Fecal samples were collected in 1200 mL 2-piece specimen collectors (Medline, USA) in the Public Health Science Division of the Fred Hutchinson Cancer Center (IRB Protocol number 5722) and transferred into an large vinyl anaerobic chamber (Coy, USA, 37°C, 5% hydrogen, 20% carbon dioxide, balanced with nitrogen) at the Institute for Systems Biology within 20 minutes of defecation. All further processing and incubation was then run inside the anaerobic chamber. 50 g of fecal material was transferred into sterile 50 oz Filter Whirl-Paks (Nasco, USA) with sterile PBS + 0.1% L-cysteine at a 1:2.5 w/v ratio and homogenized with a Stomacher Biomaster (Seward, USA) for 15 minutes. After homogenization, each sample was transferred into three sterile 250 mL serum bottles and another 2.5 parts of PBS + 0.1% L-cysteine was added to bring the final dilution to 1:5 in PBS. 87 ug/mL inulin or an equal volume of sterile PBS buffer were added to treatment or control bottles, respectively. Samples were immediately pipetted onto sterile round-bottom 2 mL 96-well plates in triplicates. Baseline samples were aliquoted into sterile 1.5 mL Eppendorf tubes and the plates were covered with Breathe-Easy films (USA Scientific Inc., USA). Plates were incubated for 7 h at 37°C and gently vortexed every hour within the chamber. Final samples were aliquoted into 1.5 mL Eppendorf tubes at the end of incubation. Baseline and 7 h samples were kept on ice and immediately processed after sampling. 500 uL of each sample were aliquoted for metagenomics and kept frozen at −80°C before and during transfer to the commercial sequencing service (Diversigen, Inc). The remaining sample was transferred to a table-top centrifuge (Fisher Scientific accuSpin, USA) and spun at 1,500 rpm for 10 minutes. The supernatant was then transferred to collection tubes kept on dry ice and transferred to the commercial metabolomics provider Metabolon, USA, for targeted SCFA quantification.

### In vitro culturing of fecal-derived microbial communities (Study C)

Homogenized fecal samples in this study again underwent anaerobic culturing at 37°C, as described above, but with a shorter culturing time of 4 hours. The slurry was diluted 2.5x in 0.1% L-cysteine PBS buffer solution. Cultures were supplemented with the dietary fibers pectin or inulin to a final concentration of 10g/L, or a sterile PBS buffer control treatment. Aliquots were taken at 0h and 4h and further processed for measurement of SCFA concentrations, which were used to estimate experimental production flux (concentration[4h] - concentration[0h]/4h). SCFA concentrations were measured using GC-FID. Briefly, the pH of the aliquots was adjusted to 2-3 with 1% aqueous sulfuric acid solution, after which they were vortexed for 10 minutes and centrifuged for 10 minutes at 10,000 rpm. 200 uL aliquots of clear supernatant were transferred to vials containing 200 uL of MeCN and 100 uL of a 0.1% v/v 2-methyl pentanoic acid solution. The resulting solutions were analyzed by GC-FID on a Perkin Elmer Clarus 500 equipped with a DB-FFAP column (30m, 0.250mm diameter, 0.25um film) and a flame ionization detector.

### In vitro culturing of fecal-derived microbial communities (Study D)

Fecal samples were collected in 1200 mL 2-piece specimen collectors (Medline, USA) in the Public Health Science Division of the Fred Hutchinson Cancer Center (IRB Protocol number 10961) and transferred into a large vinyl anaerobic chamber (Coy, USA, 37°C, 5% hydrogen, 20% carbon dioxide, balanced with nitrogen) at the Institute for Systems Biology within 30 minutes of sample receipt. All further processing and incubation was then run inside the anaerobic chamber. 30 g of fecal material was transferred into sterile 50 oz Filter Whirl-Paks (Nasco, USA) with 90 mL sterile PBS + 0.1% L-cysteine + 0.0001% resazurin and homogenized with a Stomacher Biomaster (Seward, USA) for 5 minutes. For each individual fecal sample, triplicate baseline samples of 1500uL slurry were transferred to a deep 96-well place (Fisher Scientific, USA), sealed and centrifuged at 4000rpm for 10 minutes. 300uL of the supernatant were transferred to collection tubes and immediately frozen at −80°C. An additional 1800uL of fecal slurry was transferred into a 2mL Eppendorf tube and frozen at −80°C for metagenomic shotgun sequencing. Interventions of 100uL inulin at 625mg/L, pectin at 750mg/L or PBS were transferred to in duplicate to a new deep 96-well plate, topped with 1500uL fecal slurry each, and immediately sealed with Breathe-Easy films (USA Scientific Inc., USA). Plates were incubated for 6 h at 37°C and gently vortexed every 2 hours within the chamber. After incubation, plates were immediately centrifuged at 4000rpm for 10 minutes at room temperature and 300uL of the supernatant was again transferred to collection tubes and kept at −80°C. The frozen slurry sample for metagenomic shotgun sequencing was transferred to a commercial sequencing service (Diversigen, Inc) on dry ice. The remaining supernatant samples were kept on dry ice and transferred to the commercial metabolomics provider (Metabolon, USA) for targeted SCFA quantification.

### Metagenomic sequencing and analysis

For Study A, shallow metagenomic sequencing was performed by the sequencing vendor Diversigen, USA (i.e., their BoosterShot service). In brief, DNA was extracted from the fecal slurries with the DNeasy PowerSoil Pro Kit on a QiaCube HT (Qiagen, Germany) and quantified using the Qiant-iT Picogreen dsDNA Assay (Invitrogen, USA). Library preparation was performed with a proprietary protocol based on the Nextera Library Prep kit (Illumina, USA) and the generated libraries were sequenced on a NovaSeq (Illumina, USA) with a single-end 100bp protocol. Demultiplexing was performed using Illumina BaseSpace to generate the final FASTQ files used during analysis. For Study D, DNA extraction was performed under the same protocol as Study A. Libraries for Study D were prepared with the Nextera XT Library Prep kit (Illumina, USA) and sequenced with a paired-end 2x150bp protocol on a NovaSeq 6000 (Illumina, USA) yielding at least 70M reads/sample.

Preprocessing of raw sequencing reads from Study A and D was performed using FASTP ^49^. The first 5bp on the 5’ end of each read were trimmed, and the 3’ end was trimmed using a sliding window quality filter that would trim the read as soon as the average window quality fell below 20. Reads containing ambiguous base calls or with a length of less than 15bp after trimming were removed from the analysis.

Bacterial species abundances were quantified using Kraken2 v2.0.8 and Bracken v2.2 using the Kraken2 default database which was based on Refseq release 94, retaining only those species with at least 10 assigned reads ^50,51^. The analysis pipeline can be found at https://github.com/Gibbons-Lab/pipelines/tree/master/shallow_shotgun.

### Metabolomics

Targeted metabolomics were performed using Metabolon’s high-performance liquid chromatography (HPLC)–mass spectrometry (MS) platform, as described before ^52^. In brief, fecal supernatants were thawed on ice, proteins were removed using aqueous methanol extraction, and organic solvents were removed with a TurboVap (Zymark, USA). Mass spectroscopy was performed using a Waters ACQUITY ultra-performance liquid chromatography (UPLC) and Thermo Scientific Q-Exactive high resolution/accuracy mass spectrometer interfaced with a heated electrospray ionization (HESI-II) source and an Orbitrap mass analyzer operated at 35,000 mass resolution. For targeted metabolomics ultra-pure standards of the desired short-chain fatty acids were used for absolute quantification. Fluxes for individual metabolites were estimated as the rate of change of individual metabolites during the incubation period (concentration[7h] - concentration[0h]/7h).

### Model Construction

Taxonomic abundance data inferred from 16S amplicon sequencing was summarized to the genus level (as in *in vitro* cultures, *ex vivo* study B, fiber intervention samples, and samples from the Arivale cohort), or to the species level when shotgun metagenomic sequencing was available (as in *ex vivo* studies A, C and D). Abundances were used to construct all MCMMs in this analysis using the community-scale metabolic modeling platform MICOM v0.32.5 ^21^. Models were built using the MICOM build() function with a relative abundance threshold of 0.001, omitting taxa that made up less than 0.1% relative abundance. The AGORA database (v1.03) of taxonomic reconstructions summarized to the genus level for 16S data or the species level for metagenomic sequencing data was used to collect genome-scale metabolic models for taxa present in each model. Building models at the genus level for metagenomic sequencing data was explored, but was outperformed by species level models. *In silico* media were applied to the grow() function, defining the bounds for metabolic imports by the MCMM. Medium composition varied between analyses (see *Media Construction).* Steady state growth rates and metabolic fluxes for all samples were then inferred using cooperative tradeoff flux balance analysis (ctFBA). In brief, this is a two-step optimization scheme, where the first step finds the maximal biomass production rate for the full microbial community and the second step infers taxon-specific growth rates and fluxes, while maintaining community growth within a suboptimal fraction of the theoretical maximum (i.e., the tradeoff parameter), thus balancing individual growth rates and the community-wide growth rate ^21^. All models in the manuscript used a tradeoff parameter of 0.7. This parameter value was chosen through cooperative tradeoff analysis in MICOM. Multiple tradeoff parameters were tested, and the highest parameter value (i.e. the value closest to the maximal community growth rate at 1.0) that allowed most (>90%) of taxa to grow was chosen (i.e., 0.7). Predicted growth rates from the simulation were analyzed to validate correct behavior of the models. All models were found to grow with minimum community growth rate of 0.3 h^-1^. Predicted values for export fluxes of SCFAs were collected from each MCMM using the production_rates() function, which calculates the overall production from the community that would be accessible to the colonic epithelium.

### Media Construction

Individual media were constructed based on the context of each individual analysis. For the synthetic *in vitro* cultures conducted by Clark et al. (2021), a defined medium (DM38) was used that supported growth of all taxa used in the experiments, excluding *Faecalibacterium prausnitzii.* Manually mapping each component to the Virtual Metabolic Human database, we constructed an *in silico* medium with flux bounds scaled to component concentration. All metabolites were found in the database. Using the MICOM fix_medium() function, a minimal set of metabolites necessary for all models to grow to a minimum community growth rate of 0.3 h^-1^ was added to the medium - here, only iron(III) was added (*in silico* medium available here: https://github.com/Gibbons-Lab/scfa_predictions/tree/main/media).

To mimic the medium used in *ex vivo* cultures of fecally-derived microbial communities, a carbon-stripped version of a standard European diet was used. First, a standard European diet was collected from the Virtual Metabolic Human database (www.vmh.life/#nutrition) ^53^. Components in the medium which could be imported by the host, as defined by an existing uptake reaction in the Recon3D model ^54^, were diluted to 20% of their original flux, to adjust for absorption in the small intestine^54^. Additionally, host-supplied metabolites such as mucins and bile acids were added to the medium. The medium was augmented with a minimal set of metabolites required for growth of all taxa in the model database using the complete_db_medium() function within MICOM. As most carbon sources are consumed in the body and are likely not present in high concentrations in stool, this diet was then manually stripped of carbon sources by removing metabolites identified to be carbon sources for microbes. All components in the media were then diluted to 10% of their original flux to mimic the fecal microenvironment. Residual dietary fiber not easily digested including resistant starch, dextrin and cellulose, was not removed from the medium during carbon removal. The amount of this residual fiber was scaled to the dilution factor of samples in each study prior to culturing. To simulate fiber supplementation, single fiber additions were made to the medium, either pectin, inulin or fructo-oligosaccharide (1.0 mmol/gDW*h for pectin, 10.0 mmol/gDW*h for inulin, 100 mmol/gDW*h for FOS, based on carbon content reported for each).

For *in vivo* modeling, two diets were used: a high-fiber diet containing high levels of resistant starch, and a standard European diet ^53,55^. Again, both diets were collected from the Virtual Metabolic Human database (www.vmh.life/#nutrition). Each medium was subsequently adjusted to account for absorption in the small intestine by diluting metabolite flux as described previously. Additionally, host-supplied metabolites such as mucins and bile acids were added to the medium, to match the composition of the medium *in vivo*. Finally, the complete_db_medium() function was again used to augment the medium, as described above.

Prebiotic interventions were designed by supplementing the high-fiber or average European diet with single fiber additions, either pectin or inulin, as described previously.

### Probiotic Intervention

To model a probiotic intervention, 5% relative abundance of the genus *Faecalibacterium*, a known butyrate-producing taxon ^56^, was added to the MCMMs by adding a pan-genus model of the taxon derived from the AGORA database (v1.03). Measured taxonomic abundances were scaled to 95% of their initial values, after which *Faecalibacterium* was artificially added to the model.

### External Data Collection

Data containing taxonomic abundance, optical density, and endpoint butyrate concentration for synthetically-constructed *in vitro* microbial cultures were collected from Clark et al. (2021) ^30^. Endpoint taxonomic abundance data, calculated from fractional read counts collected via 16S amplicon sequencing, was used to construct individual MCMMs for each co-culture (see *Model Construction)*. Resulting models ranged in taxonomic richness from 1 to 25 taxa.

Data from *ex vivo* studies A and D, containing shotgun metagenomic sequencing and SCFA metabolomics, was collected and processed as described previously. Taxonomic abundance data was used to construct MCMMs for each individual (see *Model Construction*).

From a study by Cantu-Jungles et al. (2021) ^18^ (*ex vivo* Study B), preprocessed taxonomic abundance and SCFA metabolomics data was collected. Homogenized fecal samples in this study underwent a similar culturing process, with a culturing time of 24 hours. Cultures were supplemented with the dietary fiber pectin, or a PBS control. Initial and endpoint metabolomic SCFA measurements were used to estimate experimental production flux (concentration[24h] - concentration[0h]/24h). Taxonomic abundance data was used to construct MCMMs for each individual.

Data from a third (Study C) was collected from the Pharmaceutical Biochemistry Group at the University of Geneva, Switzerland, under study protocol 2019-00632, containing sequencing data in FASTQ format and targeted metabolomics SCFA measurements.

Data was collected from Wastyk, et al 2021 ^31^, which provided 16S amplicon sequencing data at 9 timepoints spanning 14 weeks, along with immunological phenotyping, for 18 participants undergoing a high-fiber dietary intervention. Only 7 timepoints spanning 10 weeks were included in subsequent analysis, as the last 2 timepoints were taken after the conclusion of the dietary intervention. MCMMs were constructed for each participant at each timepoint at the genus level (see *Model Construction)*. Mean total butyrate and propionate production were compared between immune response groups.

De-identified data was obtained from a former scientific wellness program run by Arivale, Inc. (Seattle, WA) ^32^. Arivale closed its operations in 2019. Taxonomic abundances, inferred from 16S amplicon sequencing data, for 2,687 research-consenting individuals were collected and used to construct MCMMs. 128 paired blood-based clinical chemistries taken within 30 days of fecal sampling were also collected and used to find associations between MCMM SCFA predictions on a standard European diet and clinical markers. Blood pressure and BMI for each individual were also collected. Metadata for each sample including age, sex, and microbiome sequencing vendor were included in the analysis as confounders.

### Statistical analysis

Statistical analysis was performed using SciPy (v1.9.1) and statsmodels (v0.14.0) in Python (v3.8.13). Pearson correlation coefficients and p-values were calculated between measured and predicted SCFA production fluxes in *in vitro* and *ex vivo* cultures, as well as for predicted SCFA production fluxes across timepoints for an *in vivo* high-fiber intervention. Significance in SCFA production between immune response groups in the high-fiber dietary intervention was determined by non-parametric pairwise Mann-Whitney U test for butyrate, propionate, and combined butyrate and propionate production. Association of MCMM-predicted SCFA production flux with paired blood-based clinical labs was tested using OLS regression, adjusting for age, sex, microbiome sequencing vendor, and tested for significance by two-sided t-test. BMI was not included as a confounder in the analysis because it was itself negatively correlated with butyrate production ^43^. Multiple comparison correction for p-values was done using the Benjamini–Yekutieli method for adjusting the False Discovery Rate (FDR) ^57^. Comparison of butyrate production between dietary interventions was tested using paired Student’s t-tests. In all analyses, significance was considered at the p<0.05 threshold.

### Data, Software, and Code Availability

Code used to run analysis and create figures for this manuscript can be found at https://github.com/Gibbons-Lab/scfa_predictions.

Processed data for synthetically constructed cultures can be found at https://github.com/RyanLincolnClark/DesignSyntheticGutMicrobiomeAssemblyFunction. Raw sequencing data can be found at https://doi.org/10.5281/zenodo.4642238.

Raw sequencing data for Study A can be found in the NCBI SRA under accession number PRJNA937304.

Processed data for *ex vivo* Study B can be found at https://github.com/ThaisaJungles/fiber_specificity. Raw sequencing data can be found in the NCBI SRA under accession number PRJNA640404.

Raw sequencing data for *ex vivo* Study C can be found in the NCBI SRA under accession number PRJNA939256.

Raw sequencing data for *ex vivo* Study D can be found in the NCBI SRA under accession number PRJNA1033794.

Processed data for the longitudinal high-fiber intervention study can be found at https://github.com/SonnenburgLab/fiber-fermented-study/.

Qualified researchers can access the full Arivale deidentified dataset supporting the findings in this study for research purposes through signing a Data Use Agreement (DUA). Inquiries to access the data can be made at and will be responded to within 7 business days.

Illustrations were created with BioRender.com.

## Acknowledgements

We thank members of the Gibbons Lab for helpful discussions and suggestions regarding this work. Thanks to Nathan Price, Amy Willis, and Lauren Rajakovich for helpful input on this work.

## Funding

This research was funded by Washington Research Foundation Distinguished Investigator Award and by startup funds from the Institute for Systems Biology (to SMG). Fecal sample collection at Fred Hutchinson Cancer Center was supported by P30 CA015704. Research reported in this publication was supported by the National Institute of Diabetes and Digestive and Kidney Diseases of the National Institutes of Health (NIH) under award no. R01DK133468 (to SMG), by the Global Grants for Gut Health from Yakult and Nature Portfolio (to SMG), and by the National Institute on Aging of the National Institutes of Health (NIH) under award no. U19AG023122 (to NR).

## Author contributions

N.Q.B., S.M.G. and C.D. conceptualized the study. N.Q.B. ran the analyses, interpreted results and authored the first draft of the manuscript. S.M.G. and C.D. provided funding, materials and resources for the work, and supervised the work. S.M.G, C.D. and K.R.S. performed the *ex vivo* fermentation and sampling included in Study A and D. C.D. ran metagenomic analysis. J.W.L., L.L., O.V., E.M.O., K.R.S. and T.G. contributed data and resources. T.W. and N.R. provided support with analyses and statistical interpretation. All authors reviewed and edited the manuscript.

## Supplemental Figures and Captions

**Figure S1.**
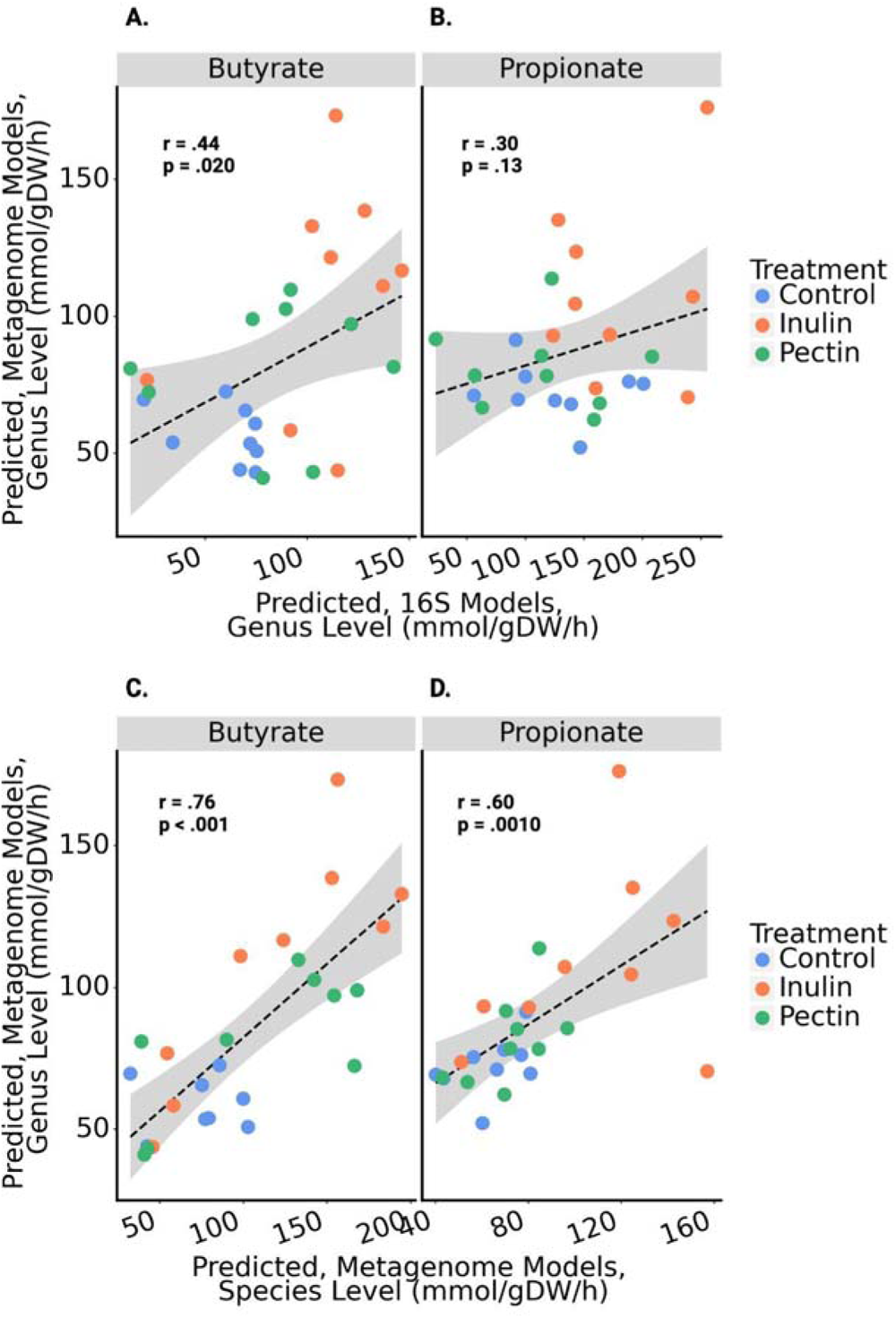
Predictions of SCFA production using 16S amplicon sequencing or shotgun metagenomic sequencing data show concordance. Data from Study C included 16S amplicon sequencing as well as shotgun metagenomic sequencing. The black line denotes a linear regression line and the gray area denotes the 95% confidence interval of the regression. Color encoding indicates the specific fiber treatment given to each sample. **(A-B)** Predictions for butyrate and propionate between models summarized to the genus level from 16S amplicon sequencing data and shotgun metagenome data. **(C-D)** Predictions for butyrate and propionate from models built using shotgun metagenome data at the genus level and species level.

**Figure S2.**
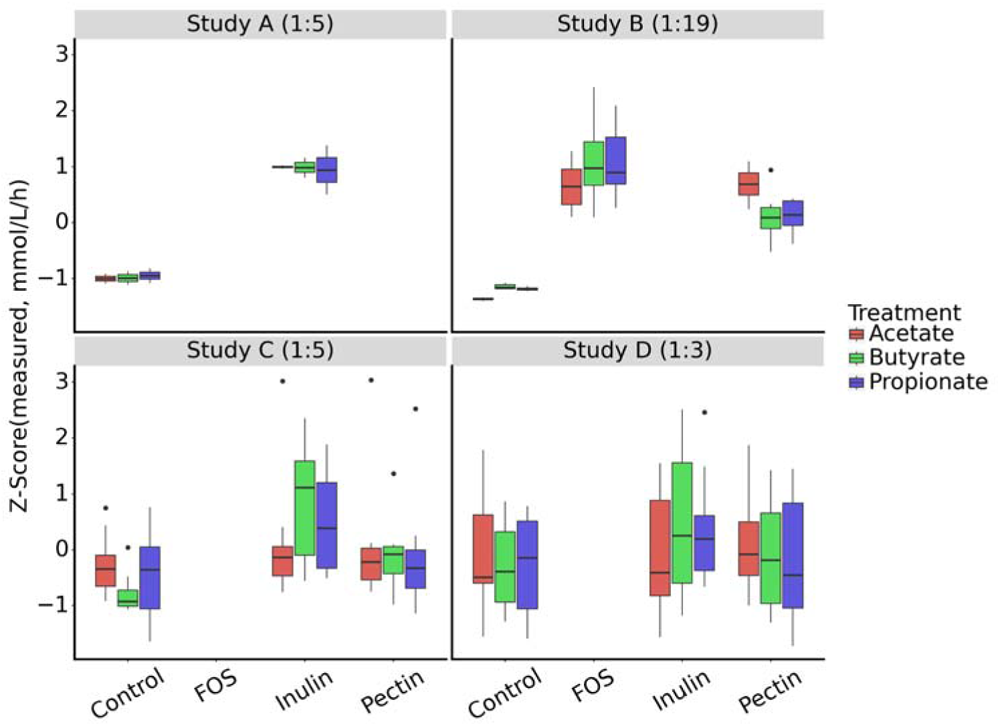
Divergence in SCFA production between controls and fiber-treated samples is related to culture dilution. Four independent *ex vivo* studies were used to validate predictions of MCMMs. Each study used a different dilution for the final culture, changing the scale of substrates available to the microbial communities. Illustrated here, the dilution factor, shown next to the study name, seems to show agreement with the divergence in SCFA production between control samples and fiber-treated samples. This was accounted for by diluting the residual fiber available to the microbial communities in the *in silico* medium.

**Figure S3.**
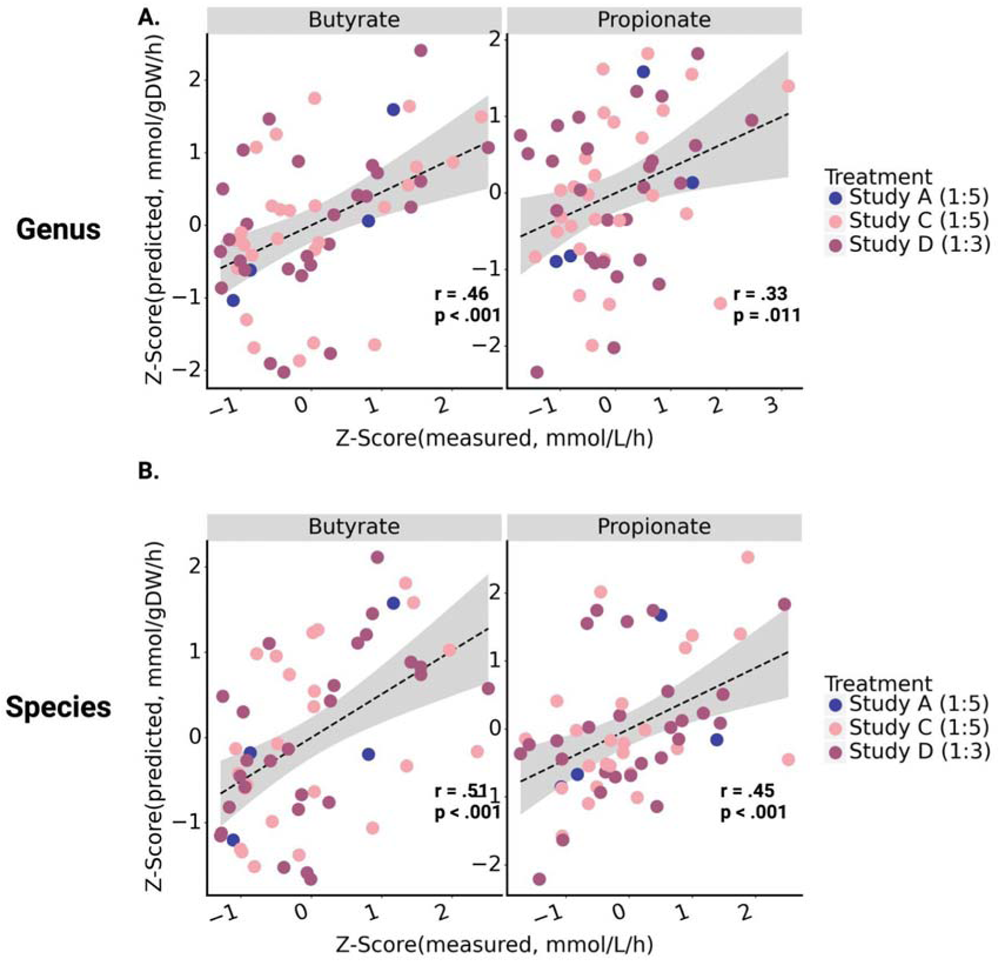
MCMMs built from shotgun metagenomic sequencing data perform better when constructed at the species level, as compared to the genus level. MCMMs from *ex vivo* studies A, C and D were constructed at the **(A)** genus and **(B)** species level. Prediction production rate of butyrate and propionate more closely matched measured production rate in the species level model as compared to the genus level model. The black line denotes a linear regression line and the gray area denotes the 95% confidence interval of the regression. Color encoding indicates the specific treatment from which Pearson r and associated p-value were calculated for each panel.

**Figure S4.**
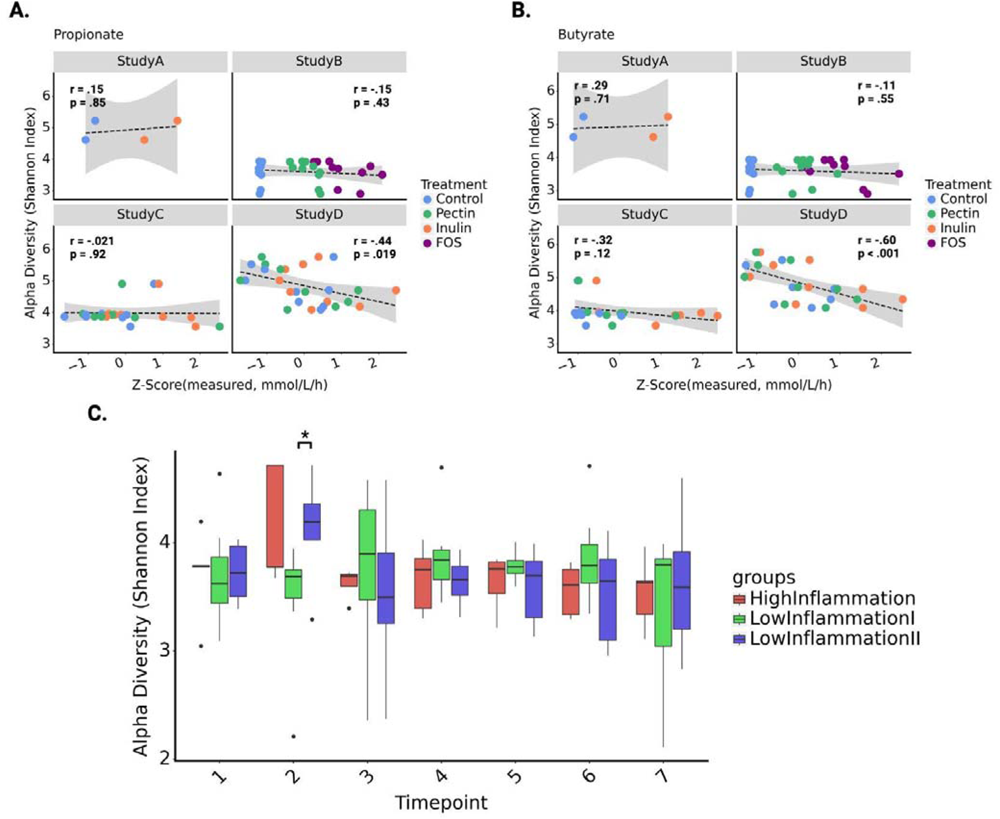
Alpha diversity of communities does not account for differences in SCFA production. We compared Shannon index, a measure of alpha diversity, against SCFA production in *ex vivo* communities, as well as between immune response groups in a longitudinal high fiber study. **(A)** Propionate production in four *ex vivo* datasets was not consistently explained by alpha diversity. In study D, a significant relationship was observed as determined by t-test (p < 0.05), but this was not consistent between datasets. The black line denotes a linear regression line and the gray area denotes the 95% confidence interval of the regression. Color encoding denotes the specific fiber treatment that was given to each sample **(B)** Butyrate production also showed no consistent correlation with alpha diversity, although a significant difference was again observed within Study D as determined by t-test (p < 0.05). **(C)** No consistent pattern emerged with regard to alpha diversity between immune response groups throughout the course of the high fiber dietary intervention, as determined by Mann Whitney U test for significance. In C, * = p < 0.05.

## Notes

https://github.com/Gibbons-Lab/scfa_predictions

https://doi.org/10.5281/zenodo.4642238

https://github.com/RyanLincolnClark/DesignSyntheticGutMicrobiomeAssemblyFunction

https://github.com/ThaisaJungles/fiber_specificity

### Competing Interest Statement

One of our coauthors, Dr. Thomas Gurry, works for a commercial prebiotics company (Myota GmbH, Inc.). Myota was not involved in the funding or conduct of this research. The authors report no other conflicts of interest.

### Summary of Updates

We have included additional analyses of MCMM performance at the genus vs. species levels, comparison of 16S vs. metagenomic data as initial constraints, and we have added an additional ex vivo data set.

## Citations

1. Oliphant, K. & Allen-Vercoe, E. Macronutrient metabolism by the human gut microbiome: major fermentation by-products and their impact on host health. Microbiome 7, 91 (2019).

2. Rackerby, B., Van De Grift, D., Kim, J. H. & Park, S. H. Effects of Diet on Human Gut Microbiome and Subsequent Influence on Host Physiology and Metabolism. Gut Microbiome and Its Impact on Health and Diseases 63–84 Preprint at 10.1007/978-3-030-47384-6_3 (2020).

3. Tomasova, L., Grman, M., Ondrias, K. & Ufnal, M. The impact of gut microbiota metabolites on cellular bioenergetics and cardiometabolic health. Nutr. Metab. 18, 72 (2021).

4. Glotfelty, L. G., Wong, A. C. & Levy, M. Small molecules, big effects: microbial metabolites in intestinal immunity. Am. J. Physiol. Gastrointest. Liver Physiol. 318, G907–G911 (2020).

5. Donia, M. S. & Fischbach, M. A. HUMAN MICROBIOTA. Small molecules from the human microbiota. Science 349, 1254766 (2015).

6. Diener, C. et al. Genome-microbiome interplay provides insight into the determinants of the human blood metabolome. Nat Metab 4, 1560–1572 (2022).

7. Ríos-Covián, D. et al. Intestinal Short Chain Fatty Acids and their Link with Diet and Human Health. Front. Microbiol. 7, 185 (2016).

8. Nogal, A., Valdes, A. M. & Menni, C. The role of short-chain fatty acids in the interplay between gut microbiota and diet in cardio-metabolic health. Gut Microbes 13, 1–24 (2021).

9. Silva, Y. P., Bernardi, A. & Frozza, R. L. The Role of Short-Chain Fatty Acids From Gut Microbiota in Gut-Brain Communication. Frontiers in Endocrinology vol. 11 Preprint at 10.3389/fendo.2020.00025 (2020).

10. Morrison, D. J. & Preston, T. Formation of short chain fatty acids by the gut microbiota and their impact on human metabolism. Gut Microbes 7, 189–200 (2016).

11. Cong, J., Zhou, P. & Zhang, R. Intestinal Microbiota-Derived Short Chain Fatty Acids in Host Health and Disease. Nutrients 14, (2022).

12. Yang, W. et al. Intestinal microbiota-derived short-chain fatty acids regulation of immune cell IL-22 production and gut immunity. Nat. Commun. 11, 4457 (2020).

13. Scheppach, W. et al. Effect of butyrate enemas on the colonic mucosa in distal ulcerative colitis. Gastroenterology 103, 51–56 (1992).

14. Tang, Y., Chen, Y., Jiang, H., Robbins, G. T. & Nie, D. G-protein-coupled receptor for short-chain fatty acids suppresses colon cancer. Int. J. Cancer 128, 847–856 (2011).

15. Singh, N. et al. Activation of Gpr109a, receptor for niacin and the commensal metabolite butyrate, suppresses colonic inflammation and carcinogenesis. Immunity 40, 128–139 (2014).

16. Tan, J. et al. The role of short-chain fatty acids in health and disease. Adv. Immunol. 121, 91–119 (2014).

17. Mortensen, P. B. & Clausen, M. R. Short-chain fatty acids in the human colon: relation to gastrointestinal health and disease. Scand. J. Gastroenterol. Suppl. 216, 132–148 (1996).

18. Cantu-Jungles, T. M. et al. Dietary Fiber Hierarchical Specificity: the Missing Link for Predictable and Strong Shifts in Gut Bacterial Communities. MBio 12, e0102821 (2021).

19. Healey, G. R., Murphy, R., Brough, L., Butts, C. A. & Coad, J. Interindividual variability in gut microbiota and host response to dietary interventions. Nutr. Rev. 75, 1059–1080 (2017).

20. Boets, E. et al. Quantification of in Vivo Colonic Short Chain Fatty Acid Production from Inulin. Nutrients 7, 8916–8929 (2015).

21. Diener, C., Gibbons, S. M. & Resendis-Antonio, O. MICOM: Metagenome-Scale Modeling To Infer Metabolic Interactions in the Gut Microbiota. mSystems 5, (2020).

22. van Deuren, T., Blaak, E. E. & Canfora, E. E. Butyrate to combat obesity and obesity-associated metabolic disorders: Current status and future implications for therapeutic use. Obes. Rev. 23, e13498 (2022).

23. Zeevi, D. et al. Personalized Nutrition by Prediction of Glycemic Responses. Cell 163, 1079–1094 (2015).

24. Rein, M. et al. Effects of personalized diets by prediction of glycemic responses on glycemic control and metabolic health in newly diagnosed T2DM: a randomized dietary intervention pilot trial. BMC Med. 20, 56 (2022).

25. Gibbons, S. M. et al. Perspective: Leveraging the Gut Microbiota to Predict Personalized Responses to Dietary, Prebiotic, and Probiotic Interventions. Adv. Nutr. 13, 1450–1461 (2022).

26. Shoaie, S. et al. Quantifying Diet-Induced Metabolic Changes of the Human Gut Microbiome. Cell Metab. 22, 320–331 (2015).

27. Heinken, A. et al. Genome-scale metabolic reconstruction of 7,302 human microorganisms for personalized medicine. Nat. Biotechnol. (2023) doi:10.1038/s41587-022-01628-0.

28. Abdill, R. J., Adamowicz, E. M. & Blekhman, R. Public human microbiome data are dominated by highly developed countries. PLoS Biol. 20, e3001536 (2022).

29. Magnúsdóttir, S. et al. Generation of genome-scale metabolic reconstructions for 773 members of the human gut microbiota. Nat. Biotechnol. 35, 81–89 (2017).

30. Clark, R. L. et al. Design of synthetic human gut microbiome assembly and butyrate production. Nat. Commun. 12, 3254 (2021).

31. Wastyk, H. C. et al. Gut-microbiota-targeted diets modulate human immune status. Cell 184, 4137–4153.e14 (2021).

32. Manor, O. et al. Health and disease markers correlate with gut microbiome composition across thousands of people. Nat. Commun. 11, 5206 (2020).

33. Quigley, E. M. M. Gut bacteria in health and disease. Gastroenterol. Hepatol. 9, 560–569 (2013).

34. Guinane, C. M. & Cotter, P. D. Role of the gut microbiota in health and chronic gastrointestinal disease: understanding a hidden metabolic organ. Therap. Adv. Gastroenterol. 6, 295–308 (2013).

35. Valgepea, K. et al. Systems biology approach reveals that overflow metabolism of acetate in Escherichia coli is triggered by carbon catabolite repression of acetyl-CoA synthetase. BMC Syst. Biol. 4, 166 (2010).

36. Wolfe, A. J. The acetate switch. Microbiol. Mol. Biol. Rev. 69, 12–50 (2005).

37. Liu, H. et al. Butyrate: A Double-Edged Sword for Health? Adv. Nutr. 9, 21–29 (2018).

38. Sze, M. A., Topçuoğlu, B. D., Lesniak, N. A., Ruffin, M. T., 4th & Schloss, P. D. Fecal Short-Chain Fatty Acids Are Not Predictive of Colonic Tumor Status and Cannot Be Predicted Based on Bacterial Community Structure. MBio 10, (2019).

39. Gut microbial metabolites lower blood pressure in patients with hypertension. Nat Cardiovasc Res 2, 18–19 (2023).

40. Amiri, P. et al. Role of Butyrate, a Gut Microbiota Derived Metabolite, in Cardiovascular Diseases: A comprehensive narrative review. Front. Pharmacol. 12, 837509 (2021).

41. Tough, I. R., Forbes, S. & Cox, H. M. Signaling of free fatty acid receptors 2 and 3 differs in colonic mucosa following selective agonism or coagonism by luminal propionate. Neurogastroenterol. Motil. 30, e13454 (2018).

42. Ulven, T. Short-chain free fatty acid receptors FFA2/GPR43 and FFA3/GPR41 as new potential therapeutic targets. Front. Endocrinol. 3, 111 (2012).

43. Coppola, S., Avagliano, C., Calignano, A. & Berni Canani, R. The Protective Role of Butyrate against Obesity and Obesity-Related Diseases. Molecules 26, (2021).

44. Babaei, P., Shoaie, S., Ji, B. & Nielsen, J. Challenges in modeling the human gut microbiome. Nat. Biotechnol. 36, 682–686 (2018).

45. Gurry, T., Nguyen, L. T. T., Yu, X. & Alm, E. J. Functional heterogeneity in the fermentation capabilities of the healthy human gut microbiota. PLoS One 16, e0254004 (2021).

46. Passi, A. et al. Genome-Scale Metabolic Modeling Enables In-Depth Understanding of Big Data. Metabolites 12, (2021).

47. Gasaly, N., de Vos, P. & Hermoso, M. A. Impact of Bacterial Metabolites on Gut Barrier Function and Host Immunity: A Focus on Bacterial Metabolism and Its Relevance for Intestinal Inflammation. Front. Immunol. 12, 658354 (2021).

48. Agus, A., Clément, K. & Sokol, H. Gut microbiota-derived metabolites as central regulators in metabolic disorders. Gut 70, 1174–1182 (2021).

49. Chen, S., Zhou, Y., Chen, Y. & Gu, J. fastp: an ultra-fast all-in-one FASTQ preprocessor. Bioinformatics 34, i884–i890 (2018).

50. Wood, D. E., Lu, J. & Langmead, B. Improved metagenomic analysis with Kraken 2. Genome Biol. 20, 257 (2019).

51. Lu, J., Breitwieser, F. P., Thielen, P. & Salzberg, S. L. Bracken: estimating species abundance in metagenomics data. PeerJ Comput. Sci. 3, e104 (2017).

52. Gauglitz, J. M. et al. Enhancing untargeted metabolomics using metadata-based source annotation. Nat. Biotechnol. 40, 1774–1779 (2022).

53. Elmadfa, I. Österreichischer Ernährungsbericht 2012. 1, (2012).

54. Brunk, E. et al. Recon3D enables a three-dimensional view of gene variation in human metabolism. Nat. Biotechnol. 36, 272–281 (2018).

55. Waldmann, A., Koschizke, J. W., Leitzmann, C. & Hahn, A. Dietary intakes and lifestyle factors of a vegan population in Germany: results from the German Vegan Study. Eur. J. Clin. Nutr. 57, 947–955 (2003).

56. Zhou, L. et al. Faecalibacterium prausnitzii Produces Butyrate to Maintain Th17/Treg Balance and to Ameliorate Colorectal Colitis by Inhibiting Histone Deacetylase 1. Inflamm. Bowel Dis. 24, 1926–1940 (2018).

57. Benjamini, Y. & Hochberg, Y. Controlling the false discovery rate: A practical and powerful approach to multiple testing. J. R. Stat. Soc. 57, 289–300 (1995).

